# cando.py: Open source software for predictive bioanalytics of large scale drug-protein-disease data

**DOI:** 10.1101/845545

**Authors:** William Mangione, Zackary Falls, Gaurav Chopra, Ram Samudrala

## Abstract

Elucidating drug-protein interactions is essential for understanding the beneficial effects of small molecule therapeutics in human disease states. Traditional drug discovery methods focus on optimizing the efficacy of a drug against a single biological target of interest. However, evidence supports the multitarget theory, i.e., drugs work by exerting their therapeutic effects via interaction with multiple biological targets. Analyzing drug interactions with a library of proteins provides further insight into disease systems while also allowing for prediction of putative therapeutics against specific indications. We present a Python package for analysis of drug-proteome and drug-disease relationships implementing the Computational Analysis of Novel Drug Opportunities (CANDO) platform [1–7]. The CANDO package allows for rapid drug similarity assessment, most notably via the bioinformatic docking protocol where billions of drug-protein interactions are rapidly scored and the similarity of drug-proteome interaction signatures is calculated. The package also implements a variety of bench-marking protocols to determine how well drugs are related to each other in terms of the indications/diseases for which they are approved. Drug predictions are generated through consensus scoring of the most similar compounds to drugs known to treat a particular indication. Support for comparing and ranking novel chemical entities, as well as machine learning modules for both benchmarking and putative drug candidate prediction is also available. The CANDO Python package is available on GitHub at https://github.com/ram-compbio/CANDO, through the Conda Python package installer, and at http://compbio.org/software/.

## Introduction

Drugs and small molecule compounds exert therapeutic effects via the perturbation of multiple macromolecules, especially proteins. Growing evidence suggests small molecule drugs interact with multiple proteins to enact cellular changes, contrary to the “magic bullet” philosophy often practiced in drug discovery [8–10]. Therefore, interpreting the totality of protein interactions for drugs provides greater insight into their therapeutic functions, with the potential for more efficient drug discovery. In addition, drug repurposing has emerged as a valuable alternative to traditional drug discovery pipelines, potentially easing the burden associated with common clinical trial failures [11–14].

We have developed the Computational Analysis of Novel Drug Opportunities (CANDO) platform for analysis of drug interactions on a proteomic scale, adhering to multitarget drug theory [1–7]. An overview of the platform is provided in Supporting Figure 1. CANDO version 2 (v2) is comprised of a library of 14,606 protein structures extracted from the Protein Data Bank, 2,162 human-approved drugs from DrugBank, and 2,178 indications/diseases from the Comparative Toxicogenomics Database (CTD), encompassing 18,709 drug-indication associations [15–17]. An additional set of 5,317 human only protein structures is also available. The platform relates small molecules based on their computed interactions with all protein structures, known as an interaction signature, then assesses a drug repurposing accuracy based on how similar drug-proteomic signatures are for those drugs approved to treat the same indications. The hypothesis is that drugs with similar interaction signatures will have similar behavior in biological systems and will therefore be useful against the same indications.

Here, we present cando.py, a Python package implementing the CANDO platform for convenient analyses of drug-protein interaction signatures with the ultimate goal of making novel putative drug candidate generation easy and accessible. The package may be used for validation of virtual screening methods for applications in drug discovery and repurposing, and for extending or developing novel drug discovery and repurposing platforms. The package reads in a matrix of pre-computed interaction scores with any number of proteins, along with a drug to indication mapping, which are then benchmarked. Compound-protein interaction signatures for novel compounds/drugs not present in our library are quickly computed and added to the matrix using our bioinformatic docking protocol, allowing for direct comparison and ranking relative to other drug signatures in the platform. The package can also read in any drug-drug similarity/distance matrix computed using any method the user desires, which may be benchmarked or used for drug-disease association prediction.

## Methods: CANDO Platform implementation

The CANDO platform is implemented in Python as a series of parallel pipelines with modules for the following major protocols (Figure 1).

**Figure 1:**
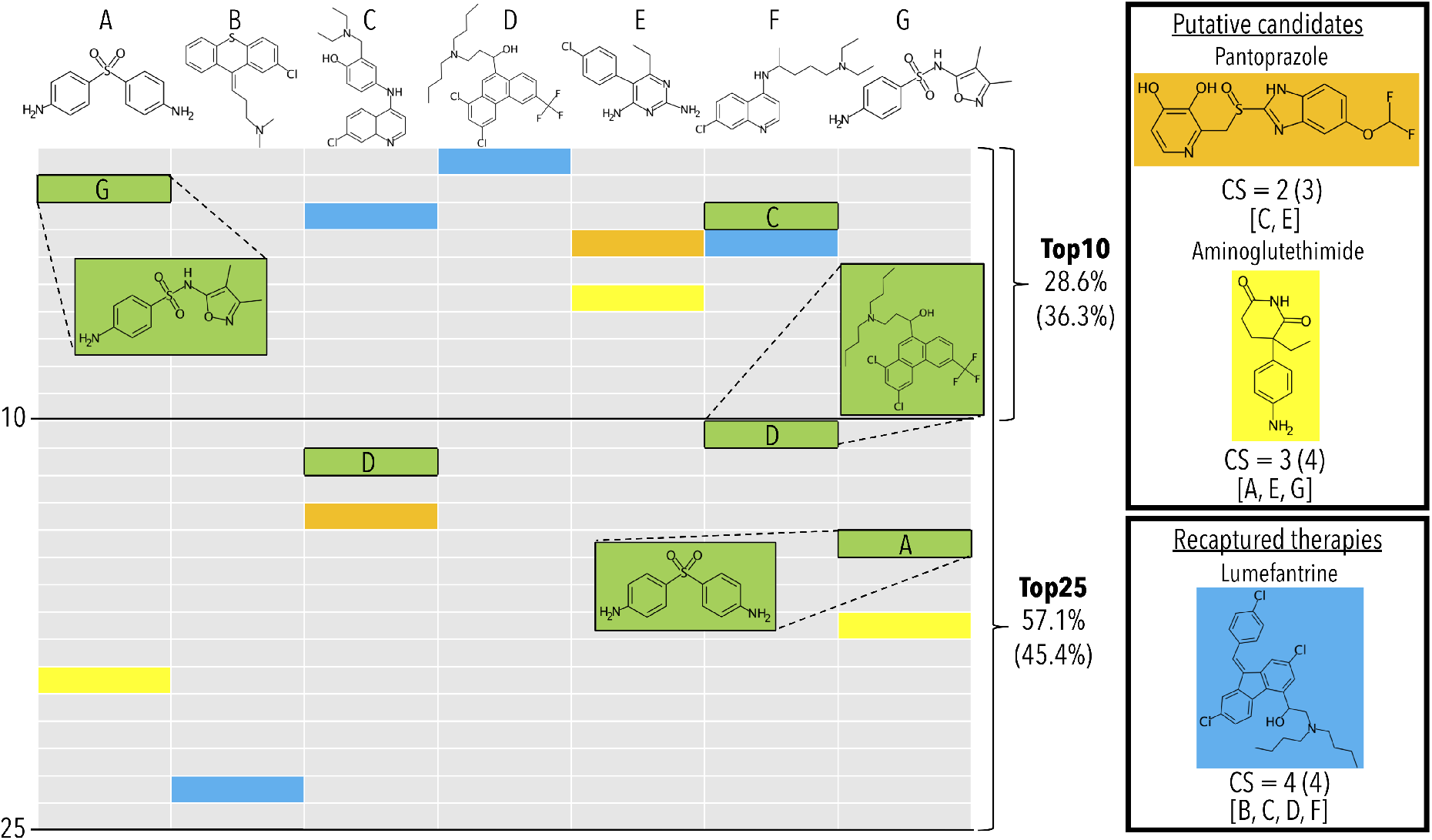
Example of benchmarking and putative therapeutic prediction with the canbenchmark and canpredict modules for malaria (Plasmodium falciparum). A subset of the 22 drugs approved for, and associated with, the indication Malaria, Falciparum (MeSH:D016778) are labeled A though G, which from left to right are dapsone, chlorprothixene, amodiaquine, halofantrine, pyrimethamine, chloroquine, and sulfisoxazole. The remaining fifteen drugs are excluded for illustrative purposes only. The benchmarking accuracies for the top10 and top25 cutoffs, and the top25 consensus scores (CS), shown in the figure are based on using only the seven drug subset (top) and for all twenty-two drugs (bottom, in parenthesis). All drugs and table cells in green are used for calculating benchmarking performance, while the yellow, blue, and orange cells are the predictions highlighted by the canpredict module. The columns represent the ranked order of the most similar drugs/compounds based on root-mean-square-deviations of their proteomic signatures to their respective A through G labeled drug. The drug-proteome interaction matrix used for this example were created from the interaction scores of 5,317 human protein structures from the Protein Data Bank with a library of 2,162 approved drugs from DrugBank. The benchmarking method tallies the percent of times another drug associated with the indication is captured within a certain column cutoff rank to a held-out compound (A – G) also associated with the indication. In the example, both drugs A and F, dapsone and chloroquine, recapture another drug associated with the indication within the top10 cutoff, which are sulfisoxazole and amodiaquine, respectively. This results in a top10 accuracy of 28.6% (two out of seven). Both amodiaquine and sulfisoxazole recapture another associated drug at the top25 cutoff, which are halofantrine and dapsone, respectively. This raises the top25 accuracy to 57.1% (four out of seven). This process is iterated over all indications to calculate global accuracies at each cutoff. The canpredict module utilizes a consensus voting scheme to suggest putative drug repurposing candidates based on the similarity of their proteomic interaction signatures to each known treatment for a disease. A tally is kept of how many times a specific drug is captured within a set cutoff to each known treatment (top25 in this example). Pantoprazole (orange), which has shown anti-malarial activity in the literature, falls at rank 14 and 4 for amodiaquine and pyrimethamine, respectively, receiving a CS of 2. The aromatase inhibitor aminoglutethimide (yellow) has a CS of 3, which we are suggesting as a novel candidate treatment for malaria. Lumefantrine, a known malaria treatment which in this case was not originally included in the Comparative Toxicogenomics Database drug-indication mapping used by the platform, receives a CS of 4. If lumefantrine was originally included as a treatment, the benchmarking scores would increase to 57.1% and 85.7% at the top10 and top25 cutoffs, respectively, which highlights the importance of drug-indication mapping veracity. The benchmarking module provides insight into how well the given drug-protein interaction scoring method is relating drugs in the context of disease, while the canpredict module suggests putative drug repurposing candidates based on drug-drug similarities.

### Interaction scoring protocol

The pipelines in the CANDO platform are agnostic to the interaction scoring protocol used: The compound-protein interaction scores in CANDO may be derived from high throughput disassociation constant studies, molecular docking simulations, and/or other quantification of structure-activity relationships [18–20]. If more than one protocol is used, then it constitutes a different pipeline within the platform. The reference/default compound-protein interaction scores in the CANDO v2 matrices are computed using a bioinformatic docking protocol that compares the structures of query drugs to all ligands known to bind to a given site on a protein [5]. Specifically, the COACH algorithm is used to elucidate potential binding sites on each query protein, which uses a consensus approach via three different complementary algorithms that consider substructure or sequence similarity to known binding sites in the PDB [21]. COACH output includes a co-crystallized ligand for each potential binding site, which can then be compared to a compound/drug of interest using chemical fingerprinting methods that binarize the presence or absence of particular molecular substructures. The maximum Tanimoto coefficient between the binary vectors of the query compound and the set of all predicted binding site ligands for a protein serves as a proxy for the binding strength. The final output is a series of interaction scores between every drug/compound and every protein structure in the corresponding libraries.

### Benchmarking protocols

Each drug/compound is ranked relative to all others based on the pairwise similarity of their proteomic signatures, calculated using the root mean square deviation (RMSD) by default, resulting in a ranked list of most similar compounds. By default, all proteins in the library are used for the RMSD calculation but their composition may be varied to allow for more specific queries, both generally or on a per indication basis, which also applies to the canpredict module (discussed below). Other distance metrics, such as cosine distance, may also be used.

The benchmarking protocol (implemented in the canbenchmark module) utilizes a hold-one-out scheme to compute an accuracy for each indication. For a given indication, each approved drug is held-out and the most similar compounds (within various cutoffs) are checked to see if they are also approved for the indication (Figure 1). This protocol is run iteratively and averaged across all indications with two or more drugs approved to provide a drug repurposing accuracy at each cutoff. Both the average indication accuracy (described above) and the pairwise accuracy (the weighted average based on the number of compounds approved for the disease) are outputted, as well as the coverage, which is the number of indications with non-zero accuracy. Benchmarking performance across different versions/pipelines is available in Supporting Figure 2.

### Putative drug candidate generation (prediction) protocol

The ranked lists of most similar compounds to each drug, other than those that are used for benchmarking, are investigated as potential novel treatments. A consensus scoring approach is utilized where for each drug associated with a specific indication, the number of times a particular drug shows up within a certain cutoff of each list is counted. The prediction module canpredict then ranks the top compounds by their consensus scores. Figure 1 provides an example with malaria (*Plasmodium falciparum*). The top consensus scoring drugs include lumefantrine, a known anti-malarial drug, and pantoprazole, a proton pump inhibitor that has shown anti-malarial activity [22]. Another strong candidate is aminoglutethimide, an aromatase inhibitor with uses including Cushing’s syndrome and various cancers. The exact set of proteins used for the drug-drug similarity calculations can be modified, e.g. specifying only *Plasmodium* proteins.

### Putative indication prediction protocol

The canpredict module can also accept a small molecule compound as input, including novel chemical entities, and suggest novel indications for which they may be useful. First the proteomic signature is computed for all proteins in the platform, then the signature is compared to all other drugs in the platform. The most similar drugs to the query compound within a specified cutoff are probed for a consensus among the diseases for which they are indicated, correlative to the disease-focused canpredict module discussed above. Figure 2 presents the results for both an approved drug, ribavirin, and an investigational compound, LMK-235. Ribavirin receives a consensus score of two at the top10 cutoff for both Breast Neoplasms (MeSH:D001943) and Leukemia, Myeloid, Acute (MeSH:D015470), which is supported by clinical trials for both diseases in which ribavirin is the primary intervention [23, 24]. The three drugs contributing to these consensus scores are gemcitabine, azacitidine, and decitabine, which are all nucleoside analog anti-cancer therapies. LMK-235 is an investigational histone deacetylase inhibitor that is yet to begin human trials. The canpredict module output with a top20 cutoff includes both Pain (MeSH:D010146) and Hypertension (MeSH:D006973), which are both supported by *in vivo* experiments [25, 26].

**Figure 2:**
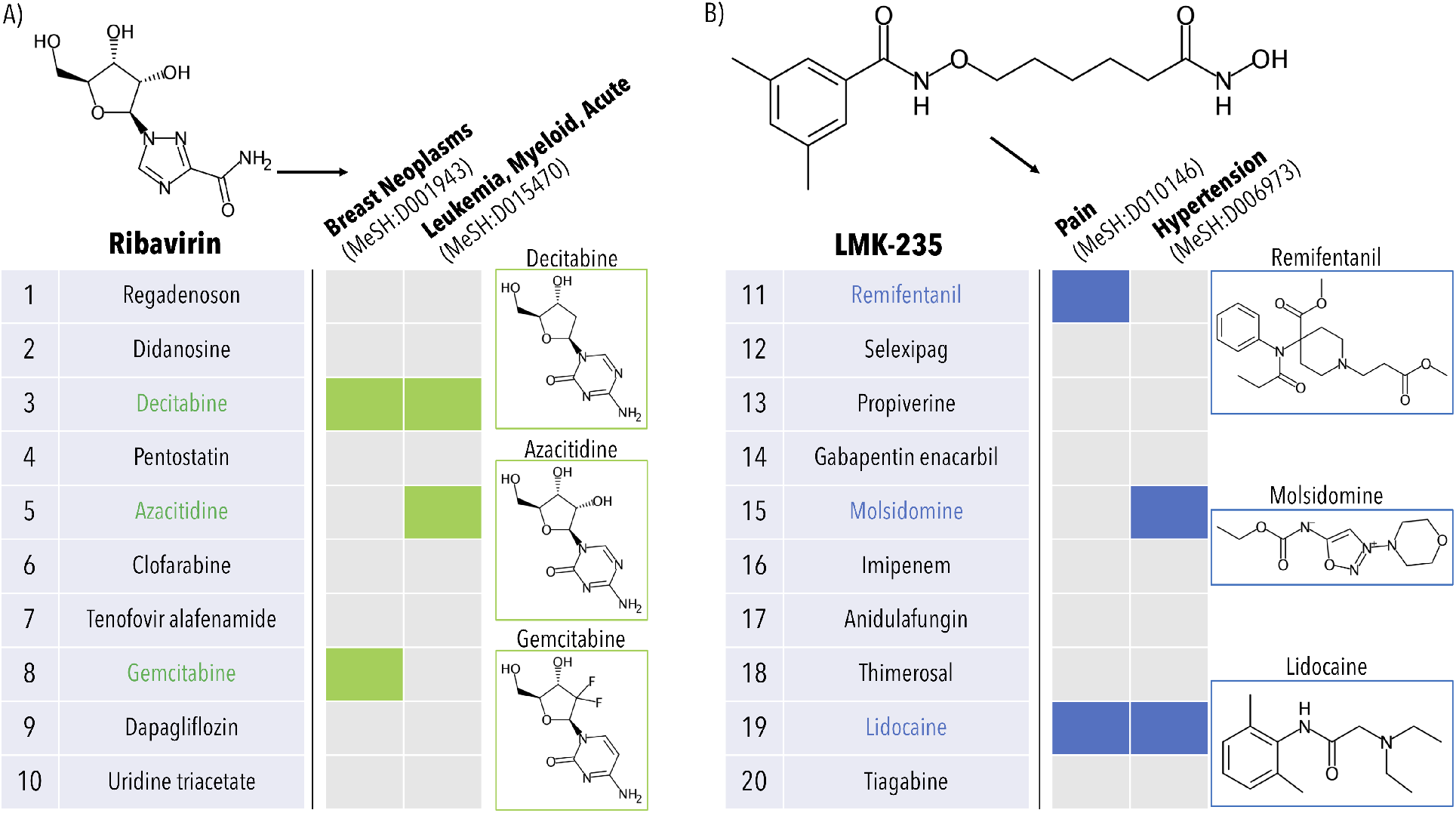
Example of indication prediction using the canpredict module with drugs (ribavirin and LMK-235). The canpredict module also accepts a drug/compound as input and suggests indications for which it may be useful based on the ranked list of the most similar drugs and the diseases for which they are approved. A) This panel presents the results for ribavirin, a known antiviral compound, using a set of 5,317 human protein structures from the Protein Data Bank to construct the drug-proteome interaction signatures. First, the top10 most similar drugs to ribavirin are computed via the root-mean-square-deviation of their proteomic interaction signatures. Then, the consensus scores of the indications associated with the top10 drugs are calculated. In the example, both Breast Neoplasms (MeSH:D001943) and Leukemia, Myeloid, Acute (MeSH:D015470) receive a consensus score of 2; these two diseases are highlighted as ribavirin is currently in clinical trials for both and has already shown clinical efficacy against acute myeloid leukemia. The three drugs contributing to the consensus scores, decitabine, azacitidine, and gemcitabine, are all chemotherapeutic nucleoside analogs. B) This panel presents the results for LMK-235, an experimental histone deacetylase inhibitor currently not approved for human use. In this case, the top20 drugs were probed for a disease consensus (only ranks 11-20 are shown for illustrative purposes). Two indications of note, namely Pain (MeSH:D010146) and Hypertension (MeSH:D006973), both receive a consensus score of 2; there exist multiple studies in the literature supporting both the analgesic and hypotensive properties of LMK-235. The drugs contributing to the consensus scoring include remifentanil, molsidomine, and lidocaine, as pictured. The CANDO canpredict module is swiftly able to assess the behavioral similarity between the known antiviral ribavirin and several antineoplastic agents, as well as between the experimental compound LMK-235 and analgesic/hypotensive drugs.

### AI/Machine learning protocols

The CANDO package also provides support for several machine learning protocols that learn more complex relationships hidden in the drug-proteome interaction signatures to improve performance. The currently supported protocols include support vector machines (SVMs), 1-class SVMs, random forests, and logistic regression, though the latter two are prioritized as they offer insight into feature importance. The modules are trained on the input data to generate models that yield prediction pipelines that are benchmarked using a protocol similar to the one used by canbenchmark: for a given indication, each approved drug is held out while the model is trained on all other drugs approved for the indication in an iterative fashion. In other words, the number of binary classifiers trained corresponds to the number of drugs associated with a particular indication. An equal number of neutral samples are chosen as negative samples during training (except 1-class SVMs), which represent drugs/compounds not associated with the indication. Several metrics are calculated based on the number of times the samples are correctly classified, including the area under the receiver operating characteristic (AUC-ROC) and precision recall (AUPR) curves (Supporting Figure 3). AUCROC and AUPR are only available with logistic regression and random forests as they offer classification probabilities. The user may also make predictions for novel or non-associated compounds after training the classifier on all approved drugs for a particular indication, and both AI/machine learning benchmarking and prediction protocols are amenable to any kind of feature input (e.g. molecular substructures instead of compound-protein interaction scores).

### Development and implementation

The CANDO software is available in Python 2.7, 3.6, and 3.7. It is available for installation via the Python Anaconda installer. All data necessary for the benchmarking and prediction modules are available for download directly in the package. The source code, API document, and a Jupyter Notebook tutorial are available on GitHub at https://github.com/ramcompbio/CANDO as well as on http://compbio.org/software/.

## Discussion and conclusion

The multitarget approach to drug discovery is vastly unexplored and shows promise for identifying novel treatments for various diseases based on the results we have obtained using our software. The CANDO Python package allows users to investigate drug-protein interactions on a proteomic scale, moving away from the single target philosophy. The multitarget approach, which in our platform is represented as the synthesis of many virtual screens, is conducive for understanding drug behavior holistically, which will allow for better elucidation of the therapeutic (and adverse) effects these small molecules exert on biological systems. We anticipate that broader use of this platform will inform researchers about potential lead compounds that may be therapeutic for specific indications, leading to accelerated and more efficient drug discovery. In addition to the bioinformatic docking protocol described above, a compound-protein interaction matrix generated using our state of the art docking program CANDOCK [27] with predicted binding energies will be available for use shortly. Indeed, the platform can accept compound-compound similarity matrices generated by any method (virtual docking, molecular fingerprinting, gene expression changes etc.). Also, a webserver hosted on compbio.org that will feature many of the functionalities described is under development.

## Acknowledgement

The authors thank Liana Bruggemann, Matthew Hudson, Manoj Mammen, and Jim Schuler for their contributions and testing of the CANDO software. This work was supported in part by a National Institute of Health Director’s Pioneer Award (DP1OD006779), a National Institute of Health Clinical and Translational Sciences Award (NCATS) (UL1TR001412), an NCATS ASPIRE design challenge award, a National Library of Medicine T15 Award (T15LM012495), a National Cancer Institute/Veterans Affairs Big Data-Scientist Training Enhancement Program Fellowship in Big Data Sciences, startup funds from the Department of Biomedical Informatics at the University at Buffalo, a start-up package from the Department of Chemistry at Purdue University, Ralph W. and Grace M. Showalter Research Trust award, the Integrative Data Science Initiative award, the Jim and Diann Robbers Cancer Research Grant for New Investigators award, and NIH NCATS ASPIRE Design Challenge awards to Gaurav Chopra. Additional support, in part by, a NCATS Clinical and Translational Sciences Award from the Indiana Clinical and Translational Sciences Institute (UL1TR002529), and the Purdue University Center for Cancer Research NIH grant P30 CA023168 are also acknowledged. The content is solely the responsibility of the authors and does not represent the official views of the National Institutes of Health.

## Supporting Information Available

The Supporting Information is available free of charge on the ACS Publications website at DOI: NUMBER.

Figure 1 - Overview of the CANDO drug discovery and repurposing platform.

Figure 2 - Benchmarking performance across multiple versions and pipelines in the CANDO platform.

Figure 3 - AI-CANDO platform performance evaluation using the receiver operating characteristic (ROC).

SMILES strings.

