## Supporting Figure 1 for "cando.py: Open source software for predictive bioanalytics of large scale drug-protein-disease data"

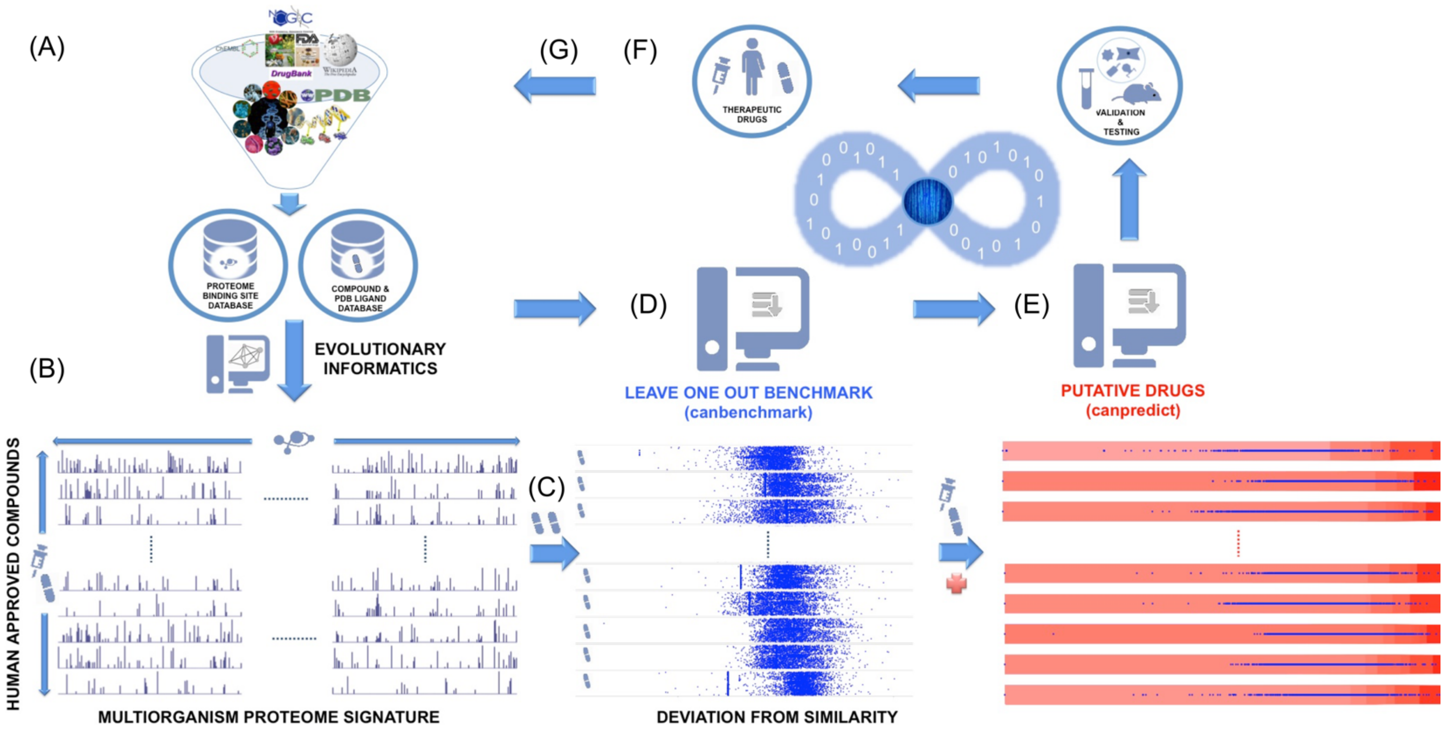


**Figure 1: Overview of the CANDO drug discovery and repurposing platform**. (A) Data collection: drugs, small molecule compounds, and protein structures are collected from public databases, most notably DrugBank and the Protein Data Bank (PDB). Different protein sets are collated, including the multiorganism sequence non-redundant PDB set (BLAST p-value cutoff 10e-7) of 14,606 chains, a human-only set of 5,317 chains, and various pathogenic proteomes. (B) Interaction scoring protocol: interactions are computed between all compounds and all proteins using our bioinformatic docking method or virtual docking simulations, which essentially results in a virtual interaction screen for every protein chain against every drug/compound in the library. The bioinformatic docking protocol generates billions of drug-protein interaction scores rapidly (see Methods). (C) Drug comparison protocol: drug-drug similarity is assessed via computing the similarity (typically by calculating the root-mean-square deviation (RMSD)) between two drug-proteome interaction signatures, which allows each drug to be ranked relative to each other (based on the composition of their interactions). The exact protein set to be considered for the RMSD calculation can be modified in various ways. (D) Benchmarking protocol: the platform is benchmarked using known drug-indication associations as a gold standard, primarily from the Comparative Toxicogenomics Database. Briefly, an accuracy is calculated based on the number of times another drug approved to treat a disease is captured within a certain rank in the similarity list of a hold-out drug known to treat the same disease. This is repeated for all indications in the platform with at least two drugs associated and averaged at various cutoffs. (E) Putative drug candidate generation protocol: putative drug candidates for a specific indication or disease are predicted based on the similarity of their proteomic interaction signatures to those of drugs known to treat that indication, and ranked via a consensus voting scheme where the top compounds are prioritized if they are highly ranked in multiple similarity lists of approved drugs for that indication. (F) Validation pipeline: strong candidates proceed to validation studies, including in vitro and in vivo experiments, with the ultimate goal of conducting clinical trials for FDA approval. (G) Platform optimization: All results from the validation studies are fed back into the platform, using machine learning to optimize performance.

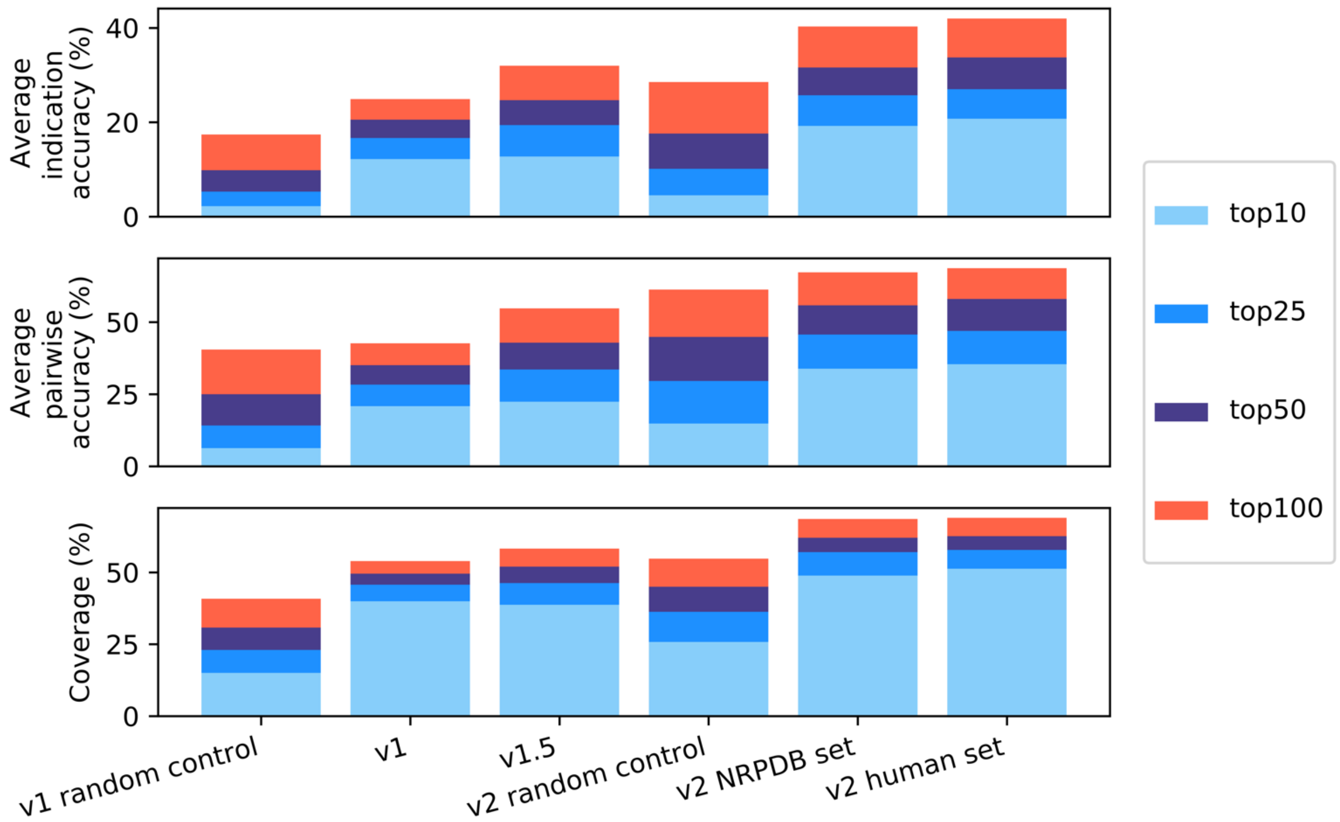


**Figure 2: Benchmarking performance across multiple versions and pipelines in the CANDO platform.** The standard three metrics assessed by the benchmarking protocol across various versions/pipelines are pictured, including average indication accuracy (top), average pairwise accuracy (middle), and coverage (bottom). Average indication accuracy is the average of each individual indication accuracy, the average pairwise accuracy is the weighted average of each indication accuracy based on the number of compounds/drugs associated with the indications, and coverage is the percent of indications with non-zero accuracy scores. The v1 and v1.5 pipelines differ only in the drug-protein interaction scoring protocol (see Falls et al. 2019), though all data including drugs, indications, and proteins is consistent. The total number of indications with greater than 2 drugs associated in v1 and v2 are 1439 and 1570, respectively. The number of drugs in v1 and v2 are 3,733 and 2,162, respectively. The number of proteins in v1 is 46,784; v2 features two protein sets: the multiorganism sequence nonredundant set of 14,606 and a set of 5,317 human protein structures, both from the Protein Data Bank. The v1 random control is computed via randomizing values in a 3,733x46,784 matrix, computing the root-mean-square-deviation between all randomized vectors, and performing the benchmarking analysis (see Methods). The result reported above is the average of 100 iterations. The v2 random control pictured was generated via a process similar to v1, but with an initial matrix of 2,162x14,606. The colors indicate using a cutoff of 10 (light blue), 25 (blue), 50 (purple), or 100 (red) for the benchmarking analysis. The results of v1 pipelines and v2 pipelines cannot be compared directly to each other as the number of drugs and drug-disease associations changes between versions. The v2 human protein set pipeline currently performs the best in terms of v2, achieving the best scores in all three metrics. The benchmarking analysis helps to indicate which pipelines are relating drugs approved for the diseases more accurately, which will ultimately help to generate novel drug repurposing candidates more efficiently.

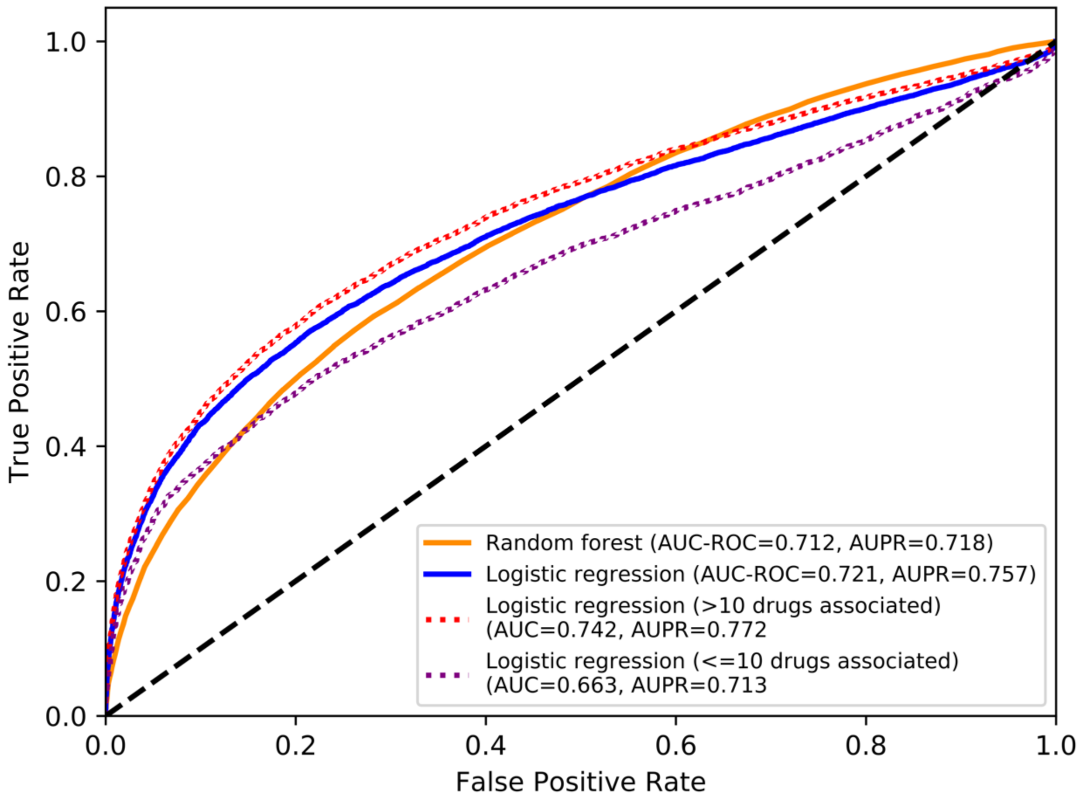


**Figure 3: AI-CANDO platform performance evaluation using the receiver operating characteristic (ROC).** Shown are the results using logistic regression and random forest binary classifiers across 1,570 indications. A leave-one-out cross validation scheme is used where each drug approved for a disease is held-out and tested on a model trained with every other drug approved for that disease (i.e. 18,101 models are trained corresponding to the number of drug-disease associations for diseases with two or more drugs associated). An equal number of negative samples are chosen by randomly selecting from the set of drugs not associated with the disease. In each case, the positive class probability is outputted and the threshold is varied to generate the receiver operating characteristic curve (ROC), area under the curve (AUC-ROC), and area under the precision-recall curve (AUPR). The dotted black line indicates the expected ROC curve if guessing randomly, which corresponds to AUC-ROC and AUPR values of 0.5. As shown, the logistic regression models (orange) outperform the random forest models (blue) in both AUC-ROC and AUPR. The AI-CANDO pipelines are based on machine learning models trained using the set of 5,317 human protein structures from the Protein Data Bank to construct the drug-proteome interaction signatures. The dotted lines indicate the difference in performance when considering indications with more than ten approved drugs (red) or with a maximum of ten approved drugs (purple), indicating the importance of sample sizes for drug-indication prediction. The model training is amenable to any input features outside of drug-protein interactions, such as molecular substructures or gene expression signatures. Though the models perform better than random, continued development of both input features (enhanced drug-protein interaction scoring) and model architecture will further improve the AI/machine learning benchmarking and prediction module.
